# Trans-ethnic genome-wide association study of kidney function provides novel insight into effector genes and causal effects on kidney-specific disease aetiologies

**DOI:** 10.1101/420273

**Authors:** Andrew P Morris, Thu H Le, Haojia Wu, Artur Akbarov, Peter J van der Most, Gibran Hemani, George Davey Smith, Anubha Mahajan, Kyle J Gaulton, Girish N Nadkarni, Adan Valladares-Salgado, Niels Wacher-Rodarte, Josyf C Mychaleckyj, Nicole D Dueker, Xiuqing Guo, Yang Hai, Jeffrey Haessler, Yoichiro Kamatani, Adrienne M Stilp, Gu Zhu, James P Cook, Johan Arnlov, Susan H Blanton, Martin H de Borst, Erwin P Bottinger, Thomas A Buchanan, Fadi J Charchar, Jeffrey Damman, James Eales, Ali G Gharavi, Vilmantas Giedraitis, Andrew C Heath, Eli Ipp, Krzysztof Kiryluk, Michiaki Kubo, Anders Larsson, Cecilia M Lindgren, Yingchang Lu, Pamela AF Madden, Holly J Mattix-Kramer, Grant W Montgomery, George J Papanicolaou, Leslie J Raffel, Ralph L Sacco, Elena Sanchez, Johan Sundstrom, Kent D Taylor, Anny H Xiang, Lars Lind, Erik Ingelsson, Nicholas G Martin, John B Whitfield, Jianwen Cai, Cathy C Laurie, Yukinori Okada, Koichi Matsuda, Charles Kooperberg, Yii-Der Ida Chen, Tanja Rundek, Stephen S Rich, Ruth JF Loos, Esteban J Parra, Miguel Cruz, Jerome I Rotter, Harold Snieder, Maciej Tomaszewski, Benjamin D Humphreys, Nora Franceschini, on behalf of the Continental Origins and Genetic Epidemiology Network (COGENT) Kidney Consortium

## Abstract

Chronic kidney disease (CKD) affects ∼10% of the global population, with considerable ethnic differences in prevalence and aetiology. We assembled genome-wide association studies (GWAS)^1-3^ of estimated glomerular filtration rate (eGFR), a measure of kidney function that defines CKD, in 312,468 individuals from four ancestry groups. We identified 93 loci (20 novel), which were delineated to 127 distinct association signals. These signals were homogenous across ancestries, and were enriched for protein-coding exons, kidney-specific histone modifications, and transcription factor binding sites for HDAC2 and EZH2. Fine-mapping revealed 40 high-confidence variants driving eGFR associations and highlighted potential causal genes with cell-type specific expression in glomerulus, and proximal and distal nephron. Mendelian randomisation (MR) supported causal effects of eGFR on overall and cause-specific CKD, kidney stone formation, diastolic blood pressure (DBP) and hypertension. These results define novel molecular mechanisms and effector genes for eGFR, offering insight into clinical outcomes and routes to CKD treatment development.

---

We assembled GWAS in up to 312,468 individuals from three sources (**Methods**): (i) 19 studies of diverse ancestry from the COGENT-Kidney Consortium, expanding the previously published trans-ethnic meta-analysis^1^ to include additional individuals of Hispanic/Latino descent; (ii) a published meta-analysis of 33 studies of European ancestry from the CKDGen Consortium^2^; and (iii) a published study of East Asian ancestry from the Biobank Japan Project^3^. Each GWAS was imputed up to the Phase 1 integrated 1000 Genomes Project reference panel^4^, and single nucleotide variants (SNVs) passing quality control were tested for association with eGFR, calculated from serum creatinine, with adjustment for age, sex and ethnicity, as appropriate (**Methods**).

To discover novel loci contributing to kidney function in diverse populations, we first aggregated eGFR association summary statistics across studies through trans-ethnic meta-analysis (**Methods**). We employed Stouffer’s method, implemented in METAL^5^, because allelic effect sizes were reported on different scales in each of the three sources contributing to the meta-analysis. We identified 93 loci attaining genome-wide significant evidence of association with eGFR (*p*<5×10^-8^), including 20 mapping outside regions previously implicated in kidney function (**Supplementary Figure 1, Supplementary Table 1**). The strongest novel associations (**Table 1**) mapped to/near *MYPN* (rs7475348, *p*=8.6×10^-19^), *SHH* (rs6971211, *p*=6.5×10^-13^), *XYLB* (rs36070911, *p*=2.3×10^-11^) and *ORC4* (rs13026220, *p*=3.1×10^-11^). Across the 93 loci, we then delineated 127 distinct association signals (at locus-wide significance, *p*<10^-5^) through approximate conditional analyses implemented in GCTA^6^ (**Methods**), each arising from different underlying causal variants and/or haplotype effects (**Supplementary Tables 1 and 2**). The most complex genetic architecture was observed at *SLC22A2* and *UMOD-PDILT*, where the eGFR association was delineated to four distinct signals at each locus (**Supplementary Figure 2**). Genome-wide, application of LD Score regression^7^ to a meta-analysis of only European ancestry studies revealed the observed scale heritability of eGFR to be 7.6%, of which 44.7%/5.4% was attributable to variation in the known/novel loci reported here (**Methods**).

**Table 1.** **Novel loci attaining genome-wide significant evidence (*p*<5×10^-8^) of association with eGFR in trans-ethnic meta-analysis of up to 312,468 individuals of diverse ancestry**.

| Locus | Lead SNV | Chr | Position<br>(bp, b37) | Alleles |  | EAF | Fixed-effects meta-analysis |  |  |  |
| --- | --- | --- | --- | --- | --- | --- | --- | --- | --- | --- |
|  |  |  |  | Effect<sup>a</sup> | Other |  | *p*-value | *N* | Beta<sup>b</sup> | SE<sup>b</sup> |
| *PMF1-BGLAP* | rs2842870 | 1 | 156,200,671 | T | C | 0.632 | $1.2 \times 10^{-8}$ | 312,468 | -0.361 | 0.094 |
| *NT5C1B-RDH14* | rs13417750 | 2 | 18,681,365 | A | G | 0.189 | $1.0 \times 10^{-8}$ | 312,468 | -0.439 | 0.108 |
| *C2orf73* | rs1527649 | 2 | 54,581,356 | C | T | 0.234 | $1.5 \times 10^{-9}$ | 311,225 | -0.413 | 0.107 |
| *ORC4* | rs13026220 | 2 | 148,586,459 | G | A | 0.366 | $3.1 \times 10^{-11}$ | 312,468 | -0.265 | 0.095 |
| *NFE2L2* | rs35955110 | 2 | 178,143,371 | C | T | 0.435 | $3.9 \times 10^{-9}$ | 312,468 | -0.353 | 0.099 |
| *XYLB* | rs36070911 | 3 | 38,498,439 | G | A | 0.528 | $2.3 \times 10^{-11}$ | 312,468 | -0.296 | 0.091 |
| *AK125311* | rs856563 | 7 | 46,723,510 | C | T | 0.750 | $5.1 \times 10^{-10}$ | 309,287 | -0.455 | 0.094 |
| *SHH* | rs6971211 | 7 | 155,664,686 | T | C | 0.417 | $6.5 \times 10^{-13}$ | 309,287 | -0.350 | 0.090 |
| *NRG1* | rs4489283 | 8 | 32,399,662 | T | C | 0.296 | $1.5 \times 10^{-8}$ | 311,632 | -0.325 | 0.094 |
| *TRIB1* | rs2001945 | 8 | 126,477,978 | C | G | 0.546 | $1.6 \times 10^{-9}$ | 312,468 | -0.264 | 0.091 |
| *DCAF12* | rs61237993 | 9 | 34,130,435 | G | A | 0.666 | $4.0 \times 10^{-8}$ | 312,465 | -0.345 | 0.122 |
| *MYPN* | rs7475348 | 10 | 69,965,177 | C | T | 0.607 | $8.6 \times 10^{-19}$ | 312,468 | -0.366 | 0.095 |
| *CYP26A1* | rs4418728 | 10 | 94,839,724 | T | G | 0.539 | $1.4 \times 10^{-8}$ | 312,468 | -0.345 | 0.092 |
| *FAM53B* | rs4962691 | 10 | 126,424,137 | T | C | 0.571 | $5.0 \times 10^{-10}$ | 312,468 | -0.291 | 0.093 |
| *RASGRP1* | rs9920185 | 15 | 39,273,575 | C | A | 0.649 | $1.0 \times 10^{-8}$ | 312,468 | -0.332 | 0.094 |
| *NFAT5* | rs11641050 | 16 | 69,622,104 | C | T | 0.697 | $2.6 \times 10^{-8}$ | 312,468 | -0.283 | 0.099 |
| *JUND-LSM4* | rs8108623 | 19 | 18,408,519 | A | C | 0.695 | $4.4 \times 10^{-8}$ | 309,634 | -0.390 | 0.108 |
| *ARFRP1* | rs1758206 | 20 | 62,336,334 | T | C | 0.082 | $2.4 \times 10^{-8}$ | 163,534 | -0.546 | 0.193 |
| *NRIP1* | rs2823139 | 21 | 16,576,783 | A | G | 0.293 | $3.7 \times 10^{-9}$ | 311,637 | -0.197 | 0.093 |
| *ATP50* | rs2834317 | 21 | 35,356,706 | A | G | 0.108 | $9.5 \times 10^{-10}$ | 312,468 | -0.475 | 0.126 |

Chr: chromosome. EAF: effect allele frequency. SE: standard error.
<sup>a</sup>Effect allele is aligned to be eGFR decreasing allele.
<sup>b</sup>Beta/SE are obtained from fixed-effects meta-analysis, with inverse variance weighting of allelic effect sizes, of up to 81,829 individuals of diverse ancestry from the COGENT-Kidney Consortium, and represent absolute decrease in eGFR per effect allele.

**Table 2.** High confidence SNVs driving eGFR associations and putative causal genes through which their effects on kidney function are mediated.

| Locus | SNV | Chr | Position (bp, b37) | *p*-value | $\pi$ | Putative causal gene | Supporting evidence |
| --- | --- | --- | --- | --- | --- | --- | --- |
| *ANXA9* | rs267738 | 1 | 150,940,625 | $1.7 \times 10^{-10}$ | 55.3% | *CERS2* | Encodes p.Glu115Ala (possibly damaging, deleterious)<sup>a</sup>. |
| *CACNA1S* | rs3850625 | 1 | 201,016,296 | $2.5 \times 10^{-9}$ | 99.0% | *CACNA1S* | Encodes p.Arg1539Cys (possibly damaging, deleterious)<sup>a</sup>. |
| *GCKR* | rs1260326 | 2 | 27,730,940 | $2.0 \times 10^{-35}$ | 86.1% | *GCKR* | Encodes p.Leu446Pro (possibly damaging, tolerated)<sup>a</sup>. |
| *C2orf73* | rs10181201 | 2 | 54,799,174 | $7.4 \times 10^{-8}$ | 60.9% | *SPTBN1* | Intronic; differential expression across kidney cell types. |
| *LRP2* | rs35472707 | 2 | 169,995,581 | $1.1 \times 10^{-6}$ | 64.3% | *LRP2* | Intronic; differential expression across kidney cell types. |
| | rs60641214 | 2 | 170,199,292 | $5.6 \times 10^{-8}$ | 64.9% | *LRP2* | Intronic; differential expression across kidney cell types. |
| *CPS1* | rs1047891 | 2 | 211,540,507 | $1.5 \times 10^{-29}$ | 98.1% | *CPS1* | Encodes p.Thr1406Asn (benign, tolerated)<sup>a</sup>. |
| *PRDM8-FGF5* | rs12509595 | 4 | 81,182,554 | $4.7 \times 10^{-16}$ | 57.1% | *FGF5* | Colocalises with lead eQTL SNV. |
| *RGS14-SLC34A1* | rs3812036 | 5 | 176,813,404 | $1.0 \times 10^{-32}$ | 65.0% | *SLC34A1* | Intronic; differential expression across kidney cell types. |
| *PIP5K1B* | rs2039424 | 9 | 71,432,174 | $1.3 \times 10^{-26}$ | 50.7% | *PIP5K1B* | Intronic; differential expression across kidney cell types. |
| *WDR37* | rs80282103 | 10 | 899,071 | $2.0 \times 10^{-18}$ | 100.0% | *LARP4B* | Intronic; differential expression across kidney cell types. |
| *MPPED2* | rs7930738 | 11 | 30,605,859 | $4.7 \times 10^{-7}$ | 51.5% | *MPPED2* | Intronic; differential expression across kidney cell types. |
| *UMOD-PDILT* | rs77924615 | 16 | 20,392,332 | $1.5 \times 10^{-54}$ | 100.0% | *UMOD* | Lead eQTL SNV; differential expression across kidney cell types. |
|  |  |  |  |  |  | *GP2* | Lead eQTL SNV; differential expression across kidney cell types. |
| *DPEP1* | rs2460449 | 16 | 89,700,747 | $4.2 \times 10^{-9}$ | 97.8% | *DPEP1* | Intronic; differential expression across kidney cell types. |
| *BCAS3* | rs9895611 | 17 | 59,456,589 | $8.9 \times 10^{-28}$ | 100.0% | *BCAS3* | Intronic; differential expression across kidney cell types. |
| | rs887258 | 17 | 59,479,580 | $2.7 \times 10^{-13}$ | 62.2% | *TBX2* | Colocalises with lead eQTL SNV. |

Chr: chromosome. $\pi$ : posterior probability of association.
<sup>a</sup>PolyPhen2/SIFT predictions.

**Figure 1.**
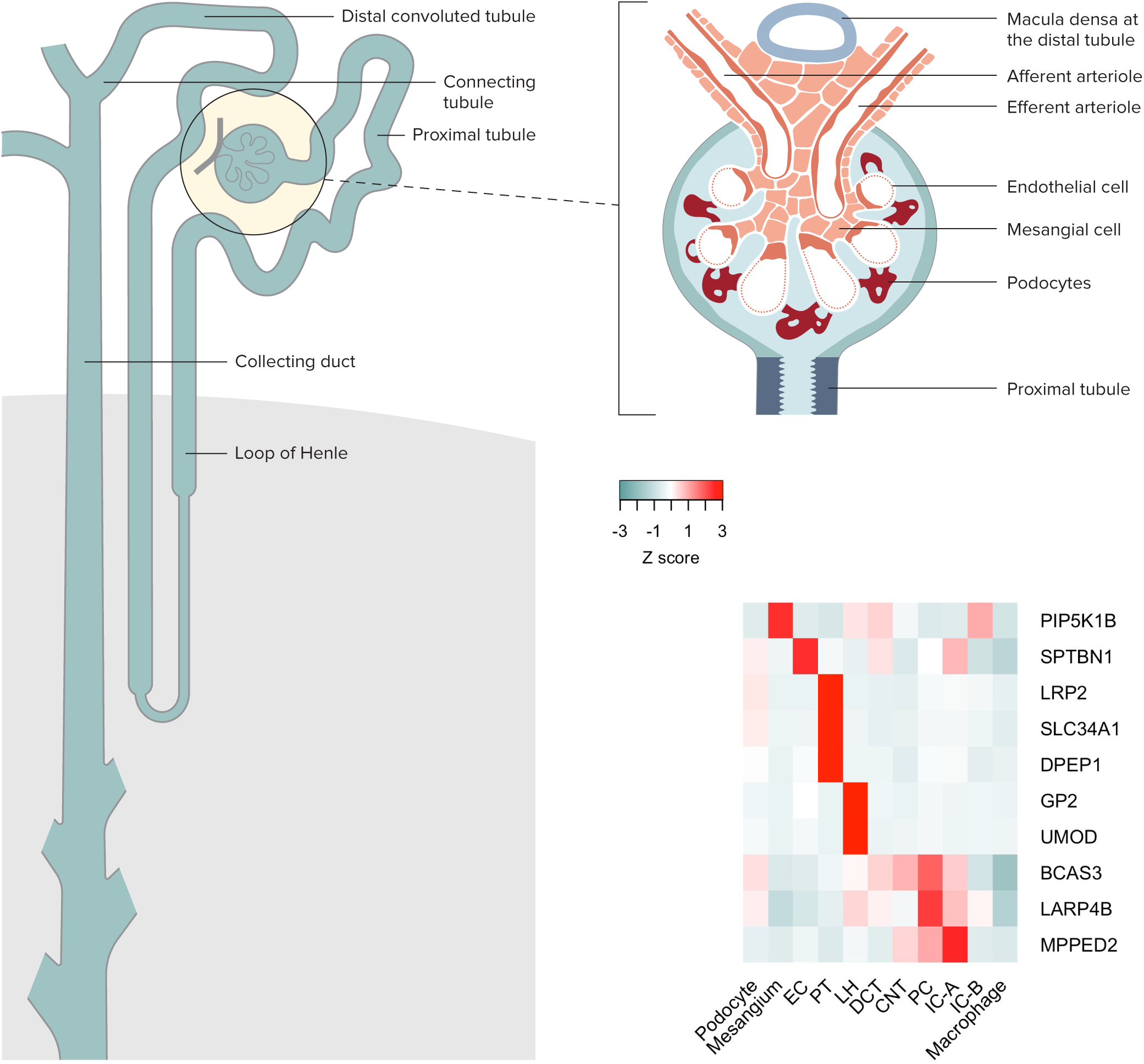
Differential kidney single cell gene expression in nephron segments. The left and top right panels highlight nephron segments and glomerulus cells, respectively. The heatmap in the bottom right panel presents *Z*-score normalized average gene expression for each specific kidney cell cluster in human adult kidney cells: EC, endothelial cells; PT, proximal tubular cells; LH, loop of Henle cells; DCT, distal convoluted cells; CNT, connecting tubular cells; PC, principal cells; IC-A, intercalate cells type A (located in the collection duct at the distal nephron); IC-B, intercalate cells type B (located in the collection duct at the distal nephron).

**Figure 2.**
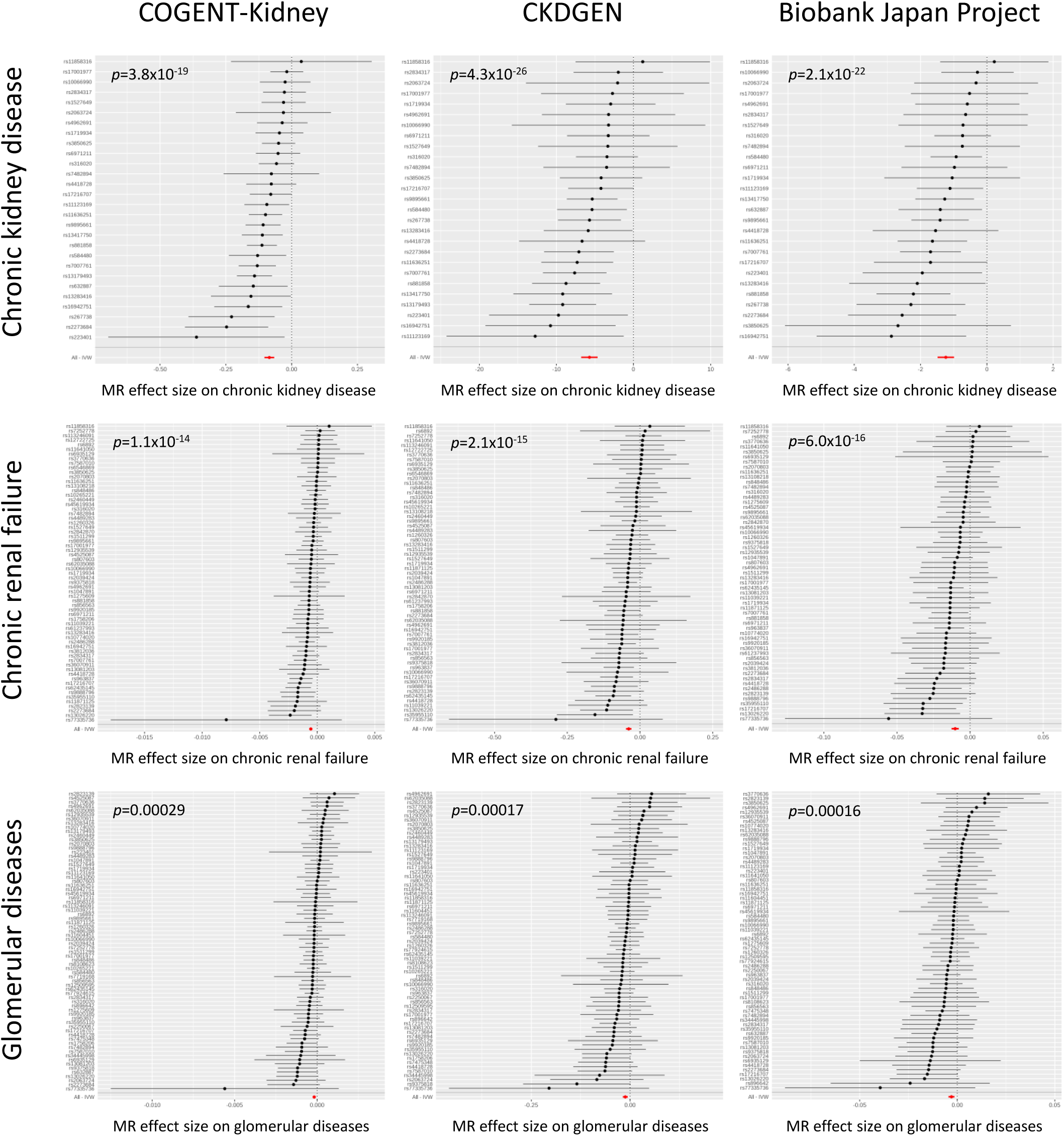
Two-sample MR of eGFR on chronic kidney disease and cause-specific kidney disease. Results are presented separately for each component of the trans-ethnic meta-analysis for chronic kidney disease (top), chronic renal failure (middle) and glomerular diseases (bottom). Each point corresponds to a lead SNV (instrumental variable) across 94 kidney function loci, plotted according to the MR effect size of eGFR on the outcome (Wald ratio). Bars correspond to the standard errors of the effect sizes. The red point and bar in each plot represents the MR effect size of eGFR on outcome across all SNVs under inverse variance weighted regression. Results for other methods are presented in **Supplementary Table 7**.

To assess the evidence for a genetic contribution to ethnic differences in CKD prevalence, we investigated differences in eGFR associations across the diverse populations contributing to our meta-analysis. We performed trans-ethnic meta-regression of allelic effect sizes obtained from GWAS contributing to the COGENT-Kidney Consortium, implemented in MR-MEGA^8^, including two axes of genetic variation that separate population groups as covariates to account for heterogeneity that is correlated with ancestry (**Methods, Supplementary Figure 3**). Despite substantial differences in allele frequencies at index SNVs for the distinct associations across ethnicities, we observed no significant evidence (*p*<0.00039, Bonferroni correction for 127 signals) of heterogeneity in allelic effects on eGFR that was correlated with ancestry (**Supplementary Tables 2 and 3**). This observation is consistent with a model in which causal variants for eGFR as a measure of kidney function are shared across global populations and arose prior to human population migration out of Africa.

**Figure 3.**
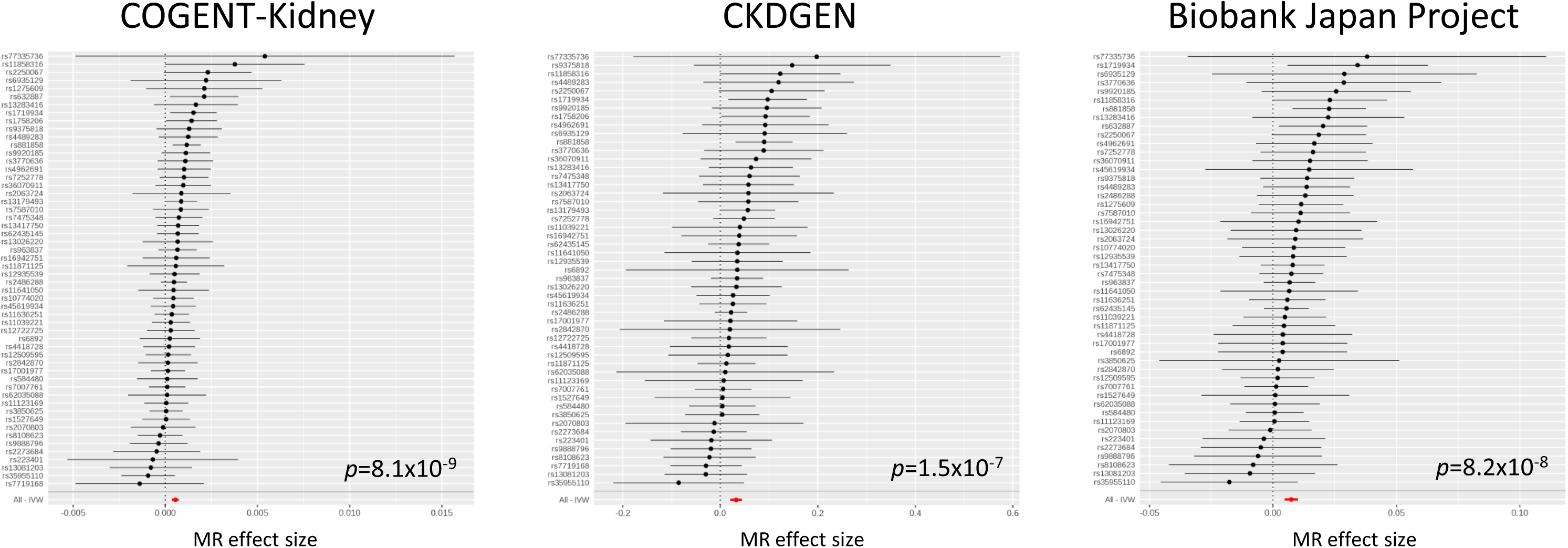
Two-sample MR of eGFR on calculus of kidney and ureter. Results are presented separately for each component of the trans-ethnic meta-analysis. Each point corresponds to a lead SNV (instrumental variable) across 94 kidney function loci, plotted according to the MR effect size of eGFR on calculus of kidney and ureter (Wald ratio). Bars correspond to the standard errors of the effect sizes. The red point and bar in each plot represents the MR effect size of eGFR on calculus of kidney and ureter across all SNVs under inverse variance weighted regression. Results for other methods are presented in **Supplementary Table 7**.

To gain insight into the molecular mechanisms that underlie the genetic contribution to kidney function, we investigated genomic signatures of functional and regulatory annotation that were enriched for eGFR associations across the 127 distinct association signals. Specifically, we compared the odds of eGFR association for SNVs mapping to each annotation with those that did not map to the annotation (**Methods**). We began by considering genic regions, as defined by the GENCODE Project^9^, and observed significant enrichment (*p*<0.05) of eGFR associations in protein-coding exons (*p*=0.0049), but not in 3’ or 5’ UTRs. We then interrogated chromatin immuno-precipitation sequence (ChIP-seq) binding sites for 161 transcription factors from the ENCODE Project^10^, which revealed significant joint enrichment of eGFR associations for HDAC2 (*p*=0.0088) and EZH2 (*p*=0.030). Class I histone deacetylases (including HDAC2) are required for embryonic kidney gene expression, growth and differentiation^11^, whilst EZH2 participates in histone methylation and transcriptional repression^12^. Finally, we considered ten groups of cell-type-specific regulatory annotations for histone modifications (H3K4me1, H3K4me3, H3K9ac and H3K27ac)^13,14^. Significant enrichment of eGFR associations was observed only for kidney-specific annotations (*p*=7.4×10^-14^). In a joint model of these four enriched annotations, the odds of eGFR association for SNVs mapping to protein-coding exons, binding sites for HDAC2 and EZH2, and kidney-specific histone modifications were increased by 3.06-, 2.13-, 1.76- and 4.29-fold, respectively (**Supplementary Figure 4**).

**Figure 4.**
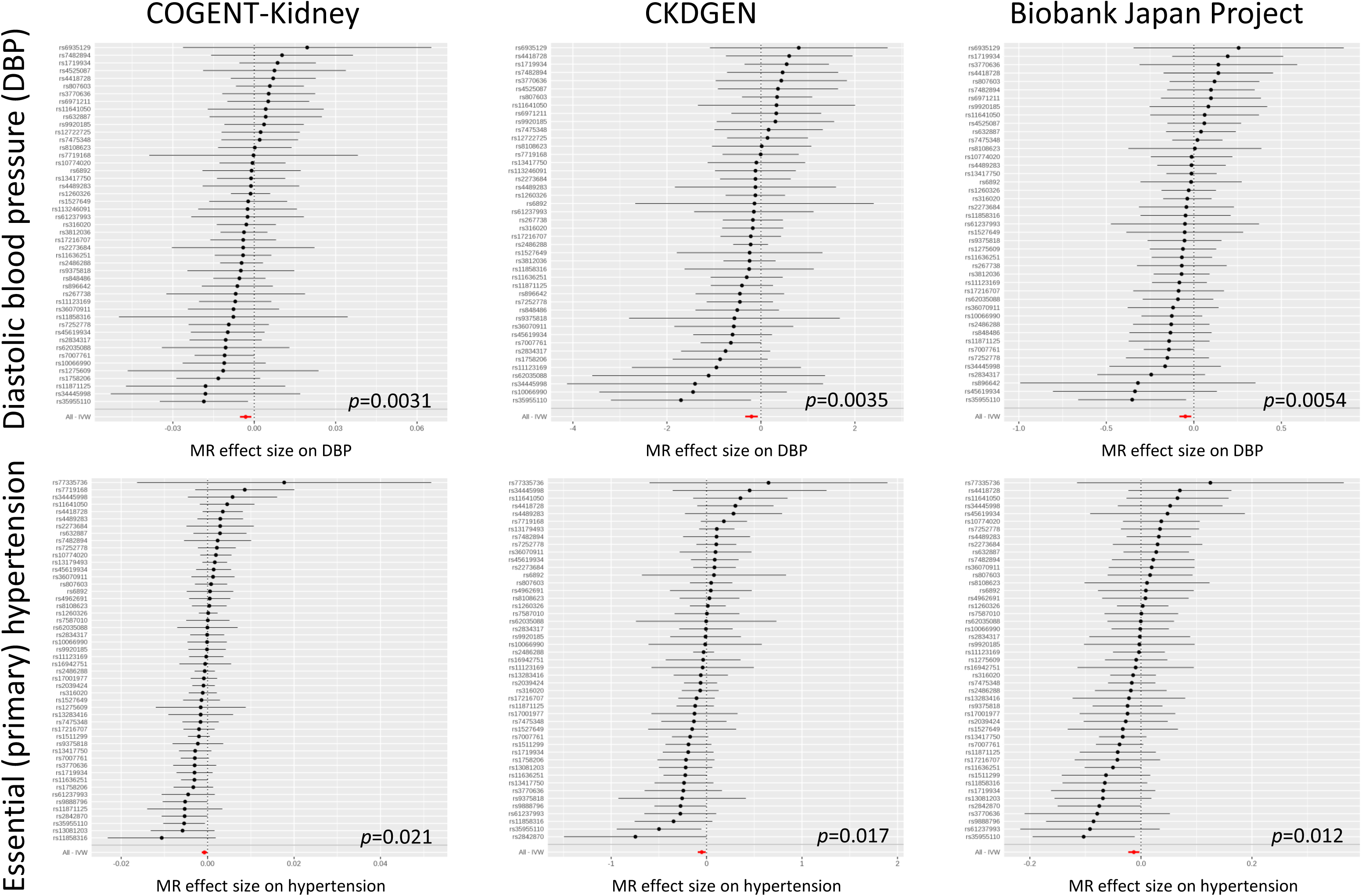
Two-sample MR of eGFR on diastolic blood pressure and essential (primary) hypertension. Results are presented separately for each component of the trans-ethnic meta-analysis for diastolic blood pressure (top) and essential (primary) hypertension (bottom). Each point corresponds to a lead SNV (instrumental variable) across 94 kidney function loci, plotted according to the MR effect size of eGFR on outcome (Wald ratio). Bars correspond to the standard errors of the effect sizes. The red point and bar in each plot represents the MR effect size of eGFR on outcome across all SNVs under inverse variance weighted regression. Results for other methods are presented in **Supplementary Table 7**.

We performed trans-ethnic fine-mapping to localise putative causal variants for distinct eGFR association signals that are shared across global populations by taking advantage of differences in the structure of linkage disequilibrium between ancestry groups^15^. To further enhance fine-mapping resolution, we incorporated an “annotation-informed” prior model for causality, upweighting SNVs mapping to the globally enriched genomic signatures of eGFR associations (**Methods**). Under this prior, we derived “credible sets” of variants for each distinct signal, which together account for 99% of the posterior probability (π) of driving the eGFR association (**Supplementary Table 4**). For 40 signals, a single SNV accounted for more than 50% of the posterior probability of driving the eGFR association, which we defined as “high-confidence” for causality (**Supplementary Table 5**).

We sought to identify the most likely target gene(s) through which the effects of each of the 40 high-confidence SNVs on eGFR were mediated. Only four of the SNVs were missense variants (**Table 2**), encoding *CACNA1S* p.Arg1539Cys (rs3850625, *p*=2.5×10^-9^, π=99.0%), *CPS1* p.Thr1406Asn (rs1047891, *p*=1.5×10^-29^, π=98.1%), *GCKR* p.Leu446Pro (rs1260326, *p*=2.0×10^-35^, π=86.1%) and *CERS2* p.Glu115Ala (rs267738, *p*=1.7×10^-10^, π=55.3%). *CACNA1S* (Calcium Voltage-Gated Channel Subunit Alpha 1 S) encodes a subunit of L-type calcium channel located within the glomerular afferent arteriole, is the target of anti-hypertensive dihydropyridine calcium channel blockers (such as amlodipine and nifedipine), and regulates arteriolar tone and intra-glomerular pressure^16^. *CACNA1S* missense mutations cause hypokalemic periodic paralysis^17,18^, malignant hyperthermia^19^ and congenital myopathy^20^. *CPS1* (Carbamoyl-Phosphate Synthase 1) is involved in the urea cycle, where the enzyme plays an important role in removing excess ammonia from cells^21^. *CERS2* (Ceramide Synthase 2) variants have been associated with albuminuria in individuals with diabetes^22^, and cers2 deficient mice exhibit changes in the structure of the kidney^23^. *GCKR* (Glucokinase Regulator) produces a regulatory protein that inhibits glucokinase, and the p.Leu446Pro substitution is a highly pleiotropic variant with reported effects on a wide range of phenotypes, including metabolic traits and type 2 diabetes^24^.

The 36 remaining high-confidence SNVs mapped to non-coding regions, which we assessed for colocalisation with expression quantitative trait loci (eQTL) from two resources: (i) non-cancer affected healthy kidney tissue obtained from 260 individuals from the TRANScriptome of renaL humAn TissuE (TRANSLATE) Study^25,26^ and The Cancer Genome Atlas (TCGA)^27^; and (ii) kidney biopsies obtained from 134 healthy donors from the TransplantLines Study^28^ (**Methods**). We observed lead eQTL variants that co-localised with high-confidence eGFR SNVs in the TRANSLATE Study and TGCA (**Table 2, Supplementary Table 6**) for *FGF5* (rs12509595, *p*=4.7×10^-16^, π=57.1%), *TBX2* (rs887258, *p*=2.7×10^-13^, π=62.2%), and both *UMOD* and *GP2* for the same signal at the *UMOD-PDILT* locus (rs77924615, *p*=1.5×10^-54^, π=100.0%). *FGF5* (Fibroblast Growth Factor 5) is expressed during kidney development, but knockout models have not shown a kidney phenotype^29^. *FGF5* has been implicated in GWAS of blood pressure and hypertension^30^, and other fibroblast growth factors are increasingly recognised as contributors to blood pressure regulation through renal mechanisms^26^. *TBX2* (T-Box 2) plays a role in defining the pronephric nephron in experimental models^31^. *UMOD* encodes uromodulin (Tamm-Horsfall protein), the most abundant urinary protein. The eGFR lowering allele at the high-confidence SNV is associated with increased *UMOD* expression (**Supplementary Figure 5**), which is consistent with previous investigations that demonstrated uromodulin overexpression in transgenic mice leads to salt-sensitive hypertension and the presence of age-dependent renal lesions^32^.

Kidney cells are highly specialised in function based on their location in nephron segments. Previous investigations in mouse and human have revealed that genes at kidney trait-related loci are expressed predominantly in a single kidney cell-type^33,34^. To provide insight into potential functional processes, we mapped the four genes identified through eQTL analyses to cell types from single nucleus RNA-sequencing (snRNA-seq) data obtained from a healthy human kidney donor (4,254 cells, with an average of 1,803 detected genes per cell)^34^. *UMOD* and *GP2* demonstrated expression that was specific to epithelial cells mapping to the ascending loop of Henle (**Figure 1**), suggesting a role for both uromodulin and glycoprotein 2, a protein involved in innate immunity, in kidney physiology at the *UMOD-PDILT* locus. By mapping high-confidence SNVs to introns and UTRs (**Methods**), we identified eight additional genes with differential expression across cell-types (**Figure 1, Table 2**): *LRP2*, *SLC34A1* and *DPEP1* (specific to proximal tubule); *SPTBN1* (specific to glomeruli endothelial cells); *PIP5K1B* (specific to glomeruli mesangial cells); and *LARP4B*, *BCAS3*, and *MPPED2* (multiple cell types in the distal nephron). Taken together, these findings suggest a potential role of these genes in influencing kidney structure and function through regulation of: (i) glomerular capillary pressure, determining intra-glomerular pressure and glomerular filtration; (ii) proximal tubular reabsorption, affecting tubuloglomerular feedback; or (iii) distal nephron handling of sodium or acid load, influencing kidney disease progression. Laboratory-based functional studies will be required to delineate the mechanistic pathways that determine kidney function in healthy and disease states, and potential routes to therapeutic targets for pharmacologic development.

We sought to evaluate the causal effect of eGFR on clinically-relevant kidney and cardiovascular outcomes via two-sample MR^35^ (**Methods, Supplementary Table 7**). Analyses were performed separately in each of the three components of the trans-ethnic meta-analysis because allelic effect sizes were measured on different scales in each. For each trait, we accounted for heterogeneity in causal effects of eGFR via radial regression^36^, excluding outlying genetic instruments that may reflect pleiotropic SNVs and violate the assumptions of MR (**Methods**). In each component, we detected a significant (*p*<0.0042, Bonferroni correction for 12 traits) causal effect of lower eGFR on higher risk of all-cause CKD, glomerular diseases and chronic renal failure, based on reported association summary statistics from the CKDGen Consortium^37^ and the UK Biobank (**Figure 2, Supplementary Table 7**). We also detected a significant causal effect of lower eGFR on lower risk of calculus of the kidney and ureter, in each component, based on reported association summary statistics from the UK Biobank (**Figure 3, Supplementary Table 7**). The lead eGFR SNV at the *UMOD-PDILT* locus (rs77924615) has been previously associated with kidney stone formation^38^ and is consistent with the role of uromodulin in the inhibition of urine calcium crystallisation^39^. However, this SNV was excluded from the MR analysis due to heterogeneity in effect size and was therefore not driving the causal eGFR association with risk of calculus of the kidney and ureter.

We also detected a novel causal effect of lower eGFR (at nominal significance, *p*<0.05, in each component of the trans-ethnic meta-analysis) on higher DBP and higher risk of essential (primary) hypertension, but not on systolic blood pressure, based on reported association summary statistics from automated readings and ICD10 codes from primary care data available in the UK Biobank (**Figure 4, Supplementary Table 7**). These results are consistent with a role for reduced functional nephron mass on increased peripheral arterial resistance^40^ and confirm previous findings from observational studies^41^. Although the causal association with DBP could not be replicated using published meta-analysis association summary statistics from the International Consortium for Blood Pressure^42^ (**Supplementary Table 8**), we note that their blood pressure measures were corrected for body-mass index (in addition to age and sex), and there was significant evidence of heterogeneity in effects of eGFR on outcome across SNVs, indicating potential pleiotropy due to collider bias, and consequently invalidating MR estimates. Despite the large sample sizes available for MR analyses from the CardiogramplusC4D Consortium^43^ and MEGASTROKE Consortium^44^, there was no significant evidence of a causal association of eGFR on cardiovascular disease outcomes: coronary heart disease, myocardial infarction or ischemic stroke (**Supplementary Table 7**).

In conclusion, we have undertaken the most comprehensive GWAS of eGFR in diverse populations, which has significantly enhanced knowledge of the genetic contribution to kidney function. Through trans-ethnic meta-analysis, we identified 20 novel loci for eGFR that explain an additional 5.3% of the genome-wide observed scale heritability. The effects of index SNVs for distinct eGFR association signals were consistent across major ancestry groups and enriched for specific signatures of genomic annotation. Annotation-informed trans-ethnic fine-mapping localised high-confidence causal variants driving 40 distinct eGFR association signals, the majority of which have not been previously reported. Through a variety of approaches, including colocalisation with eQTLs in human kidney, and identification of differential expression between human kidney cell types through snRNA-seq, these high-confidence variants implicated several putative effector genes that account for eGFR variation at kidney function loci. MR analyses of lead SNVs at kidney function loci highlighted previously unreported causal effects of lower eGFR on higher risk of primary glomerular diseases, lower risk of kidney stone formation, and higher DBP and risk of hypertension. Taken together, these results emphasize the importance of genetic studies of eGFR in diverse populations and their integration with cell-type specific kidney expression data for maximising gains in discovery and fine-mapping of kidney function loci, and offer the most promising route to treatment development for a disease with major public health impact across the globe.

## URLs

Biobank Japan Project GWAS summary statistics: http://jenger.riken.jp/en/result

CKDGen Consortium meta-analysis summary statistics: http://ckdgen.imbi.uni-freiburg.de/

METAL: https://genome.sph.umich.edu/wiki/METAL

LD Score regression: http://ldsc.broadinstitute.org/about/

MR-MEGA: https://www.geenivaramu.ee/en/tools/mr-mega

MR-BASE: http://www.mrbase.org/

GeneATLAS: http://geneatlas.roslin.ed.ac.uk/

RadialMR: https://github.com/WSpiller/RadialMR/

TwoSampleMR: https://github.com/MRCIEU/TwoSampleMR

## Acknowledgements

Thu H Le is supported by the NIH (R01-DK-113632). Gibran Hemani is supported by the Wellcome Trust (208806/Z/17/Z). George Davey Smith works in a unit supported by the MRC (R1209071-102). Ralph L Sacco and Tanja Rundek are supported by the NIH (R37-NS-029993 and U54-TR-002736) and the Evelyn F McKnight Brain Institute. Esteban J Parra is supported by the Canadian Institutes of Health Research and the Banting and Best Diabetes Center. Maciej Tomaszewski is supported the British Heart Foundation (PG/17/35/33001). Nora Franceschini is supported by the NIH (R01-MD-012765 and R56-DK-104806). Additional funding and acknowledgements can be found in the **Supplementary Note**.

## Author contributions

Central analysis: A.P.M, A.M., K.J.G. and N.F.

COGENT-Kidney Consortium GWAS analysis: A.P.M., A.M., G.N.N., A.V.-S., N.W.-R., J.C.M.,

N.D.D., X.G., Y.H., J.H., Y.K., A.M.S., G.Z., and J.P.C.

COGENT-Kidney Consortium genotyping and phenotyping: J.A., S.H.B., E.P.B., T.A.B., A.G., A.C.H., E.Ipp, M.K., A.L., C.M.L., Y.L., P.A.F.M., H.J.M-K., G.W.M., G.P., L.J.R., R.L.S., J.S., K.D.T. and A.H.X.

COGENT-Kidney Consortium principal investigator: A.P.M, L.L., E.Ingelsson, N.G.M., J.B.W., J.C., C.C.L., Y.O., K.M., C.K., Y.-D.I.C., T.R., S.S.R., R.J.F.L, E.J.P., M.C., J.I.R. and N.F.

Kidney single-cell expression data generation and analysis: H.W. and B.D.H.

TRANSLATE/TCGA data generation and eQTL analyses: A.A., J.E., F.J.C and M.T.

TransplantLines data generation and eQTL analyses: P.v.d.M., M.H.d.B., J.D. and H.S.

Mendelian randomisation analyses: A.P.M., G.H. and G.D.S.

IgA Nephropathy GWAS: E.S., A.G.G and K.K.

Manuscript preparation: A.P.M, T.H.L., H.W., A.A., M.T., B.D.H. and N.F.

COGENT-Kidney Consortium co-ordination: A.P.M and N.F.

## ONLINE METHODS

**Ethics statement.** All human research was approved by the relevant institutional review boards and conducted according to the Declaration of Helsinki. All participants provided written informed consent.

**COGENT-Kidney Consortium: study-level analyses.** Study sample characteristics for GWAS from the COGENT-Kidney Consortium, which incorporates 81,829 individuals of diverse ancestry (32.4% Hispanic/Latino, 28.8% European, 28.8% East Asian and 10.0% African American) are presented in **Supplementary Table 9**. These GWAS include those reported previously^1^ but were expanded with the addition of further studies of Hispanic/Latino ancestry to increase the diversity of represented population groups. Samples were assayed with a range of GWAS genotyping products, and quality control was undertaken within each study (**Supplementary Table 10**). Samples were excluded because of low genome-wide call rate, extreme heterozygosity, sex discordance, cryptic relatedness, and outlying ethnicity. SNVs were excluded because of low call rate across samples and extreme deviation from Hardy-Weinberg equilibrium. Non-autosomal SNVs were excluded from imputation and association analysis. Within each study, the GWAS genotype scaffold was pre-phased^45,46^ and imputed up to the Phase 1 integrated (version 3) multi-ethnic reference panel from the 1000 Genomes Project^4^ using IMPUTEv2^46,47^ or minimac^46,48^ (**Supplementary Table 10**). Imputed variants were retained for downstream association analyses if they attained IMPUTEv2 info≥0.4 or minimac *r*^2^≥0.3.

Within each study, eGFR was calculated from serum creatinine (mg/dL), with adjustment for age, sex and ethnicity, using the four variable MDRD equation^49-51^. We tested association of eGFR with each SNV in a linear regression framework, under an additive dosage model, and with adjustment for study-specific covariates to account for confounding due to population structure (**Supplementary Table 10**). For each SNV, the association *Z*-score was derived from the allelic effect estimate and corresponding standard error. *Z*-scores and standard errors were then corrected for residual population structure via genomic control^52^ where necessary (**Supplementary Table 10**).

**CKDGen Consortium: meta-analysis.** Full details of the CKDGen Consortium meta-analysis, which incorporates GWAS in 110,517 individuals of European ancestry, have been previously published^2^. Briefly, individuals were assayed with a range of GWAS genotyping products. After quality control, GWAS scaffolds were pre-phased^45,46^ and imputed^46-48^ up to the Phase 1 integrated (version 1 or version 3) multi-ethnic or European-specific reference panels from the 1000 Genomes Project^4^. Imputed variants were retained for downstream association analyses if they attained IMPUTEv2 info≥0.4 or MaCH/minimac *r*^2^≥0.4. Within each study, eGFR was calculated from serum creatinine (mg/dL), with adjustment for age and sex, using the four variable Modification of Diet in Renal Disease (MDRD) equation^49-51^. Residuals obtained after regressing ln(eGFR) on age and sex, and study-specific covariates to account for population structure where appropriate, were tested for association with each SNV in a linear regression framework, under an additive dosage model. Association summary statistics within each GWAS were corrected for residual population structure via genomic control^52^ where necessary and were subsequently aggregated across studies, under a fixed-effects model, with inverse-variance weighting of allelic effect sizes, as implemented in METAL^5^.

From the available meta-analysis summary statistics for each SNV (downloaded from http://ckdgen.imbi.uni-freiburg.de/), we derived the association *Z*-score from the ratio of the allelic effect estimate and corresponding standard error. No further correction for population structure was required by genomic control^52^: *λ*_GC_=0.977.

**Biobank Japan Project: study-level analysis.** Full details of the Biobank Japan Project GWAS, which incorporates 143,658 individuals of East Asian ancestry, have been previously published^3^. Briefly, individuals were assayed with the Illumina HumanOmniExpressExome BeadChip or a combination of the Illumina HumanOmniExpress BeadChip and the Illumina HumanExome BeadChip. After quality control, the GWAS scaffold was pre-phased with MaCH^53^ and imputed up to the Phase 1 integrated (version 3) East Asian-specific reference panel from the 1000 Genomes Project^4^ with minimac^46,48^. Imputed variants were retained for downstream association analyses if they attained minimac *r*^2^≥0.7. For each individual, eGFR was derived from serum creatinine (mg/dL) using the Japanese coefficient-modified CKD Epidemiology Collaboration (CKD-EPI) equation^54-56^, and adjusted for age, sex, ten principal components of genetic ancestry, and affection status for 47 diseases. The resulting residuals were inverse-rank normalised and tested for association with each SNV in a linear regression framework, under an additive dosage model.

From the available GWAS summary statistics for each SNV (downloaded from http://jenger.riken.jp/en/result), we derived the association *Z*-score from the ratio of the allelic effect estimate and corresponding standard error, and subsequently corrected for residual population structure by genomic control^52^: *λ*_GC_=1.252.

**Trans-ethnic meta-analysis.** We aggregated eGFR association summary statistics across the three components: COGENT-Kidney Consortium GWAS, the Biobank Japan Project GWAS and the CKDGen Consortium meta-analysis. We performed fixed-effects meta-analysis, with sample size weighting of *Z*-scores (Stouffer’s method), as implemented in METAL^5^, because allelic effect estimates were on different scales in the contributing components. The COGENT-Kidney Consortium includes a GWAS of a subset of 23,536 individuals from those contributing to the Biobank Japan Project, which was therefore excluded from the trans-ethnic meta-analysis. Consequently, a combined sample size of 312,468 individuals contributed to the trans-ethnic meta-analysis. SNVs reported in at least 50% of the combined sample size were retained for downstream interrogation. Meta-analysis association summary statistics were corrected for residual population structure via genomic control^52^: *λ*_GC_=1.113.

The current study represents a 2.2-fold increase in sample size over the largest published GWAS of kidney function^3^. Assuming homogeneous allelic effects on eGFR across populations, we have more than 80% power to detect association (*p*<5×10^-8^) with SNVs explaining at least 0.0127% of the trait variance under an additive genetic model. This corresponds to common/low-frequency SNVs with minor allele frequency (MAF) ≥5%/≥0.5% that decrease eGFR by ≥0.0366/≥0.113 standard deviations.

**Locus definition.** We first selected lead SNVs attaining genome-wide significant evidence of association (*p*<5×10^-8^) with eGFR in the trans-ethnic meta-analysis that were separated by at least 500kb. Loci were defined by the flanking genomic interval mapping 500kb up- and downstream of lead SNVs. Where loci overlapped, they were combined as a single locus, and the lead SNV with minimal *p*-value from the meta-analysis was retained.

**Dissection of association signals.** To dissect distinct eGFR association signals at loci attaining genome-wide significance in the trans-ethnic meta-analysis, we used an iterative approximate conditional approach, implemented in GCTA^6^. Each COGENT-Kidney Consortium GWAS was first assigned to an ethnic group (**Supplementary Table 9**) represented in the 1000 Genomes Project reference panel (Phase 3, October 2014 release)^57^. The Biobank Japan Project was assigned to the East Asian ethnic group, and the CKDGen Consortium meta-analysis was assigned to the European ethnic group. Haplotypes in the 1000 Genome Project panel that were specific to the assigned ethnic group were then used as a reference for LD between SNVs across loci for the GWAS in the approximate conditional analysis.

For each locus, we first applied GCTA to the study-level association summary statistics and matched LD reference for each GWAS (or the CKDGen Consortium meta-analysis). We adjusted for the “conditional set” of variants, which in the first iteration included only the lead SNV at the locus, and aggregated *Z*-scores across studies with sample size weighting (Stouffer’s method) under a fixed-effects model, as implemented in METAL^5^. The conditional meta-analysis summary statistics were corrected for residual population structure using the same genomic control adjustment^52^ as in the unconditional analysis (*λ*_GC_=1.113). If no SNVs attained locus-wide significant (*p*<10^-5^) evidence of “residual association” with eGFR, the iterative approximate conditional analysis for the locus was stopped. Otherwise, the SNV with the strongest residual association signal was added to the conditional set. This iterative process continued, at each stage adding the SNV with the strongest residual association from the meta-analysis to the conditional set, until no remaining SNVs attained locus-wide significance. Note, that at each iteration, studies with missing association summary statistics for any SNV in the conditional set were excluded from the meta-regression analysis.

For each locus including more than one SNV in the conditional set, we then dissected each distinct association signal. We again applied GCTA to the study-level association summary statistics and matched LD reference for each GWAS (or the CKDGen Consortium meta-analysis), but this time by removing each SNV, in turn, from the conditional set of variants, and adjusting for the remainder. The conditional meta-analysis summary statistics were corrected for residual population structure using the same genomic control adjustment^52^ as in the unconditional analysis (*λ*_GC_=1.113). The SNV with the strongest residual association was defined as the “index” for the signal.

**Estimation of observed scale heritability.** We used LD Score regression^7^ to assess the contribution of variation to the observed scale heritability of eGFR. LD Score regression accounts for LD between SNVs on the basis of European ancestry individuals from the 1000 Genomes Project^57^. We therefore performed fixed-effects meta-analysis, with sample size weighting of *Z*-scores (Stouffer’s method), as implemented in METAL^5^, across European ancestry studies from the COGENT-Kidney Consortium and CKDGen Consortium (134,070 individuals), and used these association summary statistics in LD Score regression. We first calculated the contribution of genome-wide variation to the observed scale heritability of eGFR. We then partitioned the genome into previously reported and novel loci attaining genome-wide significance in the trans-ethnic meta-analysis (**Supplementary Table 1**) and calculated the observed scale heritability of eGFR attributable to each.

**Estimation of allelic effect sizes at index SNVs.** Allelic effect estimates were obtained from a meta-analysis of GWAS from the COGENT-Kidney Consortium, including 81,829 individuals of diverse ancestry (**Supplementary Table 9**), because the other components applied different transformations to eGFR prior to association analysis. The meta-analysis was performed under a fixed-effects model with inverse-variance weighting of effect sizes, implemented in METAL^5^. For loci with multiple signals of association, the allelic effect of an index SNV for each GWAS, prior to meta-analysis, was estimated by application of GCTA to the study-level association summary statistics and ancestry-matched LD reference, and adjusting for the other index SNVs at the locus. The same approach was used to obtain ethnic-specific allelic effect size estimates by implementing fixed-effects meta-analysis of GWAS within each ancestry group.

**Assessment of evidence for heterogeneity in allelic effect sizes correlated with ancestry.** We considered GWAS from the COGENT-Kidney Consortium, including 81,829 individuals of diverse ancestry (**Supplementary Table 9**), because the other components applied different transformations to eGFR prior to association analysis. We constructed a distance matrix of mean effect allele frequency differences between each pair of GWAS across a subset of SNVs reported in all studies. We implemented multi-dimensional scaling of the distance matrix to obtain two principal components that define axes of genetic variation to separate GWAS from the four major ancestry groups represented in the trans-ethnic meta-analysis. For each SNV, allelic effects on eGFR across GWAS were modelled in a linear regression framework, incorporating the three axes of genetic variation as covariates, and weighted by the inverse of the variance of the effect estimates, implemented in MR-MEGA^8^. Within this modelling framework, heterogeneity in allelic effects on eGFR between GWAS is partitioned into two components. The first component is correlated with ancestry and is accounted for in the meta-regression by the axes of genetic variation, whilst the second is the residual, which is not due to population genetic differences between GWAS.

**Enrichment of eGFR association signals in genomic annotations.** Within each locus, for each distinct signal, we first approximated the Bayes’ factor^58^ in favour of eGFR association of each SNV on the basis of summary statistics from the trans-ethnic meta-analysis. Specifically, the Bayes’ factor for the *j*th SNV at the *i*th distinct association signal is approximated by

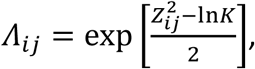

where *Z*_*ij*_ is the *Z*-score from the trans-ethnic meta-analysis across *K* contributing GWAS. The log-odds of association of the SNV is then given by

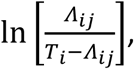

where *T*_*i*_ = ∑_*j*_ *Λ*_*ij*_ is the total Bayes’ factor for the *i*th signal across all SNVs at the locus.

We modelled the log-odds of association of each SNV, for each distinct signal, in a logistic regression framework, as a function of binary variables indicating overlap with a given genomic annotation. Specifically, for the *j*th SNV at the *i*th distinct association signal,

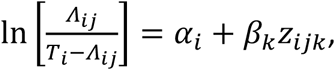

where *z*_*ijk*_ = 1 indicates that the SNV maps to the *k*th annotation, and *z*_*ijk*_ = 0 otherwise. In this expression, *α*_*i*_ is a constant for the *i*th distinct association signal, and *β*_*k*_ is the log-fold enrichment in the odds to association for the *k*th annotation.

We considered three categories of functional and regulatory annotations. First, we considered genic regions, as defined by the GENCODE Project^9^, including protein-coding exons, and 3’ and 5’ UTRs as different annotations. Second, we considered chromatin immuno-precipitation sequence (ChIP-seq) binding sites for 161 transcription factors from the ENCODE Project^10^. Third, we considered ten groups of cell-type-specific regulatory annotations for histone modifications (H3K4me1, H3K4me3, H3K9ac, and H3K27ac) obtained from a variety of resources^13,14^, which were previously derived for partitioning heritability by annotation by LD score regression^59^.

Within each category, we first used forward selection to identify annotations that were jointly enriched at nominal significance (*p*<0.05). We then included all selected annotations across categories in a final model to obtain joint estimates of the fold-enrichment in eGFR association signals for each.

**Trans-ethnic fine-mapping.** Within each locus, for each distinct signal, we calculated the posterior probability of driving the eGFR association for each SNV under an annotation-informed prior model, derived from the globally enriched annotations identified above, and the Bayes’ factor approximated from the trans-ethnic meta-analysis. Specifically, for the *j*th SNV at the *i*th distinct association signal, the posterior probability *π*_*ij*_ ∝ *γ*_*ij*_*Λ*_*ij*_. In this expression, the relative annotation informed prior is given by

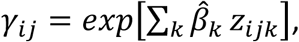

where the summation is over the selected enriched annotations, and 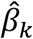 is the estimated log-fold enrichment of the *k*th annotation from the final joint model, as described above.

We derived a 99% credible set^60^ for the *i*th distinct association signal by: (i) ranking all SNVs according to their posterior probability *π*_*ij*_; and (ii) including ranked SNVs until their cumulative posterior probability of driving the association attains or exceeds 0.99. Index SNVs accounting for more than 50% posterior probability of driving the eGFR association at a given signal were defined as “high-confidence”.

**Colocalisation of high-confidence SNVs with eQTLs in kidney tissue: TRANSLATE Study and TGCA.** We performed eQTL analysis using data from the TRANSLATE Study^25,26^ and TGCA^27^. In brief, as a source of kidney tissue, both studies used apparently normal samples from European ancestry individuals undergoing nephrectomy due to kidney cancer (the specimens were collected from cancer-unaffected pole of the organ). The data from both studies were processed in the same manner using procedures described below.

Gene expression was quantified in transcripts per million (TPM) using Kallisto^61^. The quality control included: removing outlier samples^62,63^, checking consistency between declared and biological sex (using XIST and Y-chromosome genes); removing genes on non-autosomal chromosomes; and removing genes with either interquartile range of zero or those not meeting the minimum expression criterion (TPM>0.1 and read counts ≥6 in at least 30% of samples within each study/sequencing batch). Before *cis*-eQTL analysis, the log_2_-transformed TPM data were normalised using robust quantile normalisation in the R package aroma and then standardised using rank-based inverse normal transformation in GenABEL. To account for technical variation, we used probabilistic estimation of expression residuals (PEER)^64^: 30 latent factors for the TRANSLATE Study and 15 for TCGA as recommended for different sample sizes in the GTEx Project^65,66^.

Kidney DNA samples from individuals from the TRANSLATE Study were genotyped using the Infinium HumanCoreExome-24 BeadChip array, and genotype calls were made using Genome Studio. Individuals from TCGA were genotyped using the Affymetrix Genome-Wide Human SNP Array 6.0, and genotype calls were made using the Birdseed algorithm. Quality control removed variants that: had low genotyping rate (<95%); mapped to Y chromosome/mitochondrial DNA or had ambiguous chromosomal location; violated Hardy-Weinberg equilibrium (HWE, *p*<0.001); or had MAF <5%. Quality control also removed individuals with: genotyping call-rate <95%; heterozygosity above/below 3 standard deviations from the mean; cryptic relatedness to other individuals; non-European genetic ancestry; and discordant sex information (inconsistency between declared and genotyped sex). For both studies, the resulting scaffold was imputed up to the Phase 3 multi-ethnic reference panel from the 1000 Genomes Project^57^ using the Michigan Imputation Server^67^. After imputation, we retained only SNVs, removing those with low imputation coefficient (*R*^2^<0.4), MAF <5%, or violating HWE (*p*<10^-6^).

A total of 260 individuals (160 from the TRANSLATE Study and 100 from TCGA) were included in the analysis, involving 15,711 genes and 5,498,156 SNVs common to both studies. Normalised gene expression was modelled as a function of alternate allele dosage via linear regression, including sex, three axes of genetic variation (to account for population structure) and PEER latent factors as additional covariates. The regression coefficients of the alternate allele from the two studies were then combined in a fixed-effects meta-analysis under an inverse variance weighting scheme. For each gene, only those SNVs in *cis* (within 1Mb of the transcription start/stop sites) were included in the analysis. A total of 2,000 permutations were used to derive the empirical distribution of the smallest *p*-value for each gene, which then was used to adjust the observed smallest *p*-value for the gene. The correction for testing multiple genes was based on false discovery rate (FDR) applied to permutation-adjusted *p*-values (via Storey’s method as implemented in the R package qvalue) with a cut-off of 5%. Furthermore, the thresholds for nominal *p*-values were derived using a global permutation-adjusted *p*-value closest to FDR of 5% and the empirical distributions determined using permutations.

We identified high-confidence SNVs from the trans-ethnic fine-mapping that were colocalised with lead eQTL variants (i.e. the same SNV or in strong LD, *r*^2^>0.8) at a 5% FDR, and reported the corresponding eGene.

**Colocalisation of high-confidence SNVs with eQTLs in kidney tissue: TransplantLines Study.** We performed eQTL analysis using data from the TransplantLines Study^28^. The study includes kidneys from donors, donated after brain death or cardiac death. Samples were genotyped on the Illumina CytoSNP 12 v2 array and imputed up to the Phase 1 integrated (version 3) multi-ethnic reference panel from the 1000 Genomes Project^4^ using IMPUTEv2^46,47^. Expression and genotype data were available for 236 kidney biopsies obtained from 134 donors, and analyses have been described previously^42^. Briefly, residuals of gene expression for each probe were obtained after adjusting for the first 50 expression principal components to filter out environmental variation^68^. A linear mixed model was used to test association of residual expression of each probe with the allele dosage of each SNV mapping within 1Mb of the transcription start/stop sites using the R package lme3. Sex, age, donor type, time of biopsy and three axes of genetic variation (to account for population structure) were included in the model as fixed effects. Random effects were then included for donor to account for multiple samples obtained from the same individual.

We identified high-confidence SNVs from the trans-ethnic fine-mapping that were colocalised with lead eQTL variants (i.e. the same SNV or in strong LD, *r*^2^>0.8) at a 5% FDR, and reported the corresponding eGene.

**Differential expression of GWAS genes across kidney cell-types.** We identified genes for which high-confidence SNVs mapped to introns and untranslated regions. We mapped the genes to cell-types from snRNA-seq data generated by 10× Chromium from a healthy human kidney (62-year old white male, no history of CKD and serum creatinine of 1.03mg/dl)^34^. The dataset included 4,524 cells, with an average of 1,803 detected genes per cell. We generated a differential expression gene (DEG) list by performing Wilcoxon rank sum test on each cell-type from the single nucleus dataset. A gene was defined as mapping to a specific kidney cell type if the expression fulfils all the following criteria: (i) present in the DEG list; (ii) expressed in >25% of the total cells in the specified cell-type; and (iii) log-fold change in expression was >0.25 in the specified cell-type when compared to all other cell-types^34^. Gene expression values for each cell were *Z*-score normalised. A new gene expression matrix with mean *Z*-scores for each gene was obtained by averaging the *Z*-scores from all individual cells in the same cluster. The *Z*-score normalized gene expression were presented as a heatmap using the heatmap.2 function in the R package gplots.

**Two-sample MR analyses.** We performed a lookup of association summary statistics for lead SNVs at each of the eGFR loci across a range of clinically-relevant kidney and cardiovascular outcomes from public and proprietary data resources. These included: CKD (12,385 cases and 104,780 controls, published data from the CKDGen Consortium^37^); IgA nephropathy (3,211 cases and 8,735 controls, unpublished data); glomerular diseases (ICD10 N00-N08, 2,289 cases and 449,975 controls, extracted UK Biobank using GeneATLAS); chronic renal failure (ICD10 N18, 4,905 cases and 447,359 controls, extracted from UK Biobank using GeneATLAS); hypertensive renal disease (ICD10 I12, 1,663 cases and 450,601 controls, extracted from UK Biobank using GeneATLAS); calculus of kidney and ureter (ICD10 N20, 5,216 cases and 447,048 controls, extracted from UK Biobank using GeneATLAS); DBP (317,756 individuals, automated reading, extracted from UK Biobank using MR-BASE^69^); systolic blood pressure (317,654 individuals, automated reading, extracted from UK Biobank using MR-BASE^69^); essential (primary) hypertension (ICD10 I10, 84,640 cases and 367,624 controls, extracted from UK Biobank using GeneATLAS); coronary heart disease (60,801 cases and 123,504 controls, published data from the CardiogramplusC4D Consortium^43^); myocardial infarction (43,676 cases and 128,199 controls, published data from the CardiogramplusC4D Consortium^43^); and ischemic stroke (10,307 cases and 19,326 controls, published data from the MEGASTROKE Consortium^44^).

We performed two-sample MR for each outcome using eGFR as the exposure and the extracted non-palindromic lead SNVs as instrumental variables. The lead SNVs were not in LD with each other, so that their effects on exposure and outcomes were uncorrelated. Analyses were performed separately in each of the three components of the trans-ethnic meta-analysis because allelic effect sizes were measured on different scales in each: COGENT-Kidney Consortium (58,293 individuals after excluding those from the Biobank Japan Project); CDKGen Consortium (110,517 individuals); and Biobank Japan Project (143,658 individuals). For each trait, we first accounted for heterogeneity in causal effects of eGFR via radial regression^36^, implemented in the R package RadialMR, which identified outlying genetic instruments that may reflect pleiotropic SNVs. For each trait, our primary MR analyses were then performed after excluding outlying SNVs in any component of the trans-ethnic meta-analysis using inverse variance weighted regression^70^, implemented in the R package TwoSampleMR^69^. We also assessed the evidence for causal association between exposure and outcome using two additional approaches that are less sensitive to heterogeneity (although less powerful) and implemented in the R package TwoSampleMR^69^: weighted median regression^71^ and MR-EGGER regression^72^.

We performed an additional lookup of association summary statistics for non-outlying lead SNVs at each of the eGFR loci for DBP (150,134 individuals, published data from ICBP^42^). We assessed the evidence for a causal association of eGFR on DBP in each component of the trans-ethnic meta-analysis using inverse variance weighted regression^70^, weighted median regression^71^ and MR-EGGER regression^72^, as implemented in the R package TwoSampleMR^69^.

## URLs

aroma: http://aroma-project.org/

GenABEL: http://genabel.org/

Birdseed: https://www.broadinstitute.org/birdsuite/birdsuite

qvalue: https://github.com/StoreyLab/qvalue

